# Cryo-electron tomography reveals lineage-specific replication features of Zika virus

**DOI:** 10.1101/2025.08.15.670067

**Authors:** Yen-Chi Chiu, Jean Daraspe, Marco P. Alves, Christel Genoud, David Baud, Miloš Stojanov

**Affiliations:** Materno-fetal and Obstetrics Research Unit, Mother-woman-child Department, University Hospital of Lausanne, Lausanne, Switzerland; Electron Microscopy Facility, Faculty of Biology and Medicine, University of Lausanne, Lausanne, Switzerland; Institute of Virology and Immunology, Bern, Switzerland; Department of Infectious Diseases and Pathobiology, Vetsuisse Faculty, University of Bern, Bern, Switzerland; Multidisciplinary Center for Infectious Diseases, University of Bern, Bern, Switzerland; Institute of Bioengineering, School of Life Science, EPFL, Lausanne, Switzerland; Faculty of Biology and Medicine, University of Lausanne, Lausanne, Switzerland

**Keywords:** Flavivirus, Zika virus, cryo-EM, cryo-ET, Convoluted membranes, Replication complexes

## Abstract

Zika virus (ZIKV), a member of the *Flavivirus* genus, can cause severe neurodevelopmental disorders when infection occurs during pregnancy. Phylogenetic analyses classify ZIKV into two major lineages: African and Asian. Despite their genetic similarity, only the Asian lineage has been linked to congenital anomalies. ZIKV possesses a single-stranded RNA genome and replicates by remodelling the host endoplasmic reticulum (ER) to form membrane-bound replication complexes, which support viral RNA synthesis, virion assembly, and immune evasion. ZIKV also induces the formation of convoluted membranes (CMs), whose function remains unclear. Most ultrastructural studies of ZIKV replication have relied on transmission electron microscopy, which, despite its value, can alter membrane architecture due to sample preparation. To overcome these limitations, we employed cryo-focused ion beam milling combined with electron tomography to visualize ZIKV replication in a near-native state. By comparing infections with African and Asian ZIKV lineages, we uncovered lineage-specific ultrastructural features, despite similar replication dynamics and growth curves. In cells infected with the Asian strain, virions at different maturation stages were often co-packaged within the same compartment. In contrast, the African lineage ZIKV showed distinct compartmentalization of virions and vesicles. Furthermore, the ER-virion spacing was uniform in African-lineage-infected cells, while morphological heterogeneity was evident in the Asian lineage, including irregular viral sacs. Vesicle diameter was also significantly larger in cells infected with the Asian lineage. Together, these findings reveal both conserved and distinct replication structures between ZIKV lineages. The Asian lineage induces more extensive membrane remodelling and CM formation, potentially contributing to its association with congenital disease.

## Introduction

Zika virus (ZIKV), an arbovirus of the *Flavivirus* genus (recently renamed *Orthoflavivirus*), caused a major outbreak in the Americas between 2015 and 2016, which was associated not only with serious congenital defects in newborns, but also with widespread neurological complications and societal disruption, prompting the World Health Organization to declare it a Public Health Emergency of International Concern (PHEIC)^1^. The virus was first identified in 1947 in Uganda, but it remained largely neglected for decades, as it reached Asia^2^. In the 2000s, ZIKV began to emerge, eventually causing a major outbreak in 2007 on the Yap Islands in the Federated States of Micronesia^3^. A subsequent outbreak occurred in French Polynesia between 2013 and 2014, during which ZIKV infection was associated with neurological complications, including Guillain-Barré syndrome^4^. The virus then re-emerged on a much larger scale in Brazil and rapidly spread throughout Latin America between 2015 and 2016. This unprecedented epidemic led to the identification of ZIKV as the causative agent of congenital Zika syndrome (CZS), a spectrum of birth defects that includes severe brain abnormalities, such as microcephaly^5,6^. Phylogenetic analyses have classified ZIKV into two major clades, African and Asian lineages^7–9^. While being closely related, only the Asian lineage has been implicated in the abovementioned outbreaks and fetal anomalies^10,11^.

ZIKV has a single-stranded positive RNA genome encoding three structural (envelope – E, precursor-membrane – prM and capsid – C) and seven non-structural (NS1, N2A, N2B, N3, N4A, N4B and N5) proteins essential for viral replication^12^. The architecture of ZIKV-infected cells has been investigated by transmission electron microscopy (TEM)^13–16^. Consistent with other flaviviruses, ZIKV remodels the host endoplasmic reticulum (ER) to form membrane-bound replication complexes^17^, which facilitate viral RNA synthesis and virion assembly while protecting viral components from host immune responses^18,19,14^. Electron tomography (ET) has demonstrated the dynamic communication between replication vesicles and the cytoplasm, allowing the export of viral RNA to assembly sites. Additionally, ZIKV induces significant rearrangements of intracellular membranes, giving rise to convoluted membranes (CMs). Although their exact function is still unknown, studies on other flaviviruses suggest that CMs may serve as hubs for polyprotein processing^20^, or as temporary reservoirs for proteins and membranes required for viral replication^18^. Interestingly, these structures were absent in the ZIKV-infected C6/36 mosquito cell line, which did not undergo virus-induced cell death^14,21^.

The reconstruction of ZIKV particles have been carried out using cryo-electron microscopy (cryo-EM)^22–24^. A nascent virus is assembled in the ER as an immature particle, which is made of 60 trimeric E-prM heterodimer spikes. During this stage, the viral particle is non-infectious, until the exposure to low pH environment in Golgi apparatus. Simultaneously, prM protein is cleaved into the pr peptide and M protein by furin protease. These maturation steps produce smooth-surfaced ZIKV particles of approximately 50 nm^22,24^ in diameter, each composed of 90 E–M protein heterodimers, compared with the 60 nm^23^, structurally distinct immature virions.

Most of the studies on ultrastructural analysis of ZIKV replication have been performed using conventional TEM. Although these results have been invaluable for delineating ZIKV ultrastructure, the fixation and dehydration steps intrinsic to its sample preparation can artifactually perturb membrane integrity and compromise visualization of critical ultrastructural features essential to viral replication^23^. Cryo-EM has revolutionized ultrastructural biology by enabling the study of biological structures in near-native states^24–27^. Unlike conventional TEM, cryo-EM uses cryofixation by rapid freezing (vitrification) to prevent water crystallization and eliminates the need for chemical fixation and dehydration, thereby reducing artifacts^28^. To address the need for examining large cellular structures and generating thin lamellae, cryo-focused ion beam (cryo-FIB) milling and cryo-electron tomography (cryo-ET)^29,30^ have been developed in cellular biology^31–33^, enabling thinning of vitreous samples. Therefore, the combined applications preserve cellular and molecular details, offering high-resolution imaging of biological samples in their natural configuration, making it particularly valuable for examining delicate features such as membranes^34^.

In this study, we employed both conventional TEM and cryo-EM to investigate the replication dynamics and cellular impact of ZIKV infection. We utilized the human glioblastoma-derived U-87 MG cell line, a relevant model given its susceptibility to ZIKV^13,14^. By combining cryo-focused ion beam scanning electron microscopy (cryo-FIB-SEM) with cryo-ET, we visualized the ultrastructure of ZIKV replication complexes in a near-native state. Furthermore, by comparing infections with the Asian and African lineages of ZIKV, we sought to identify lineage-specific features that may influence infection outcomes.

## Materials and Methods

### Cell line and virus infection

U-87 MG cell line (ATTC HTB-14, human brain glioblastoma) and Vero cell line (ATTC CCL-81, *Cercopithecus aethiops* epithelial cells) were grown in Dulbecco’s Modified Eagle Medium (DMEM, Gibco, Basel, Switzerland) containing 10% (v/v) heat-inactivated fetal bovine serum (FBS, Sigma-Aldrich, Buchs, Switzerland) and incubated at 37°C with 5% CO_2_. ZIKV strains used in this study were the prototypic African lineage strain MP1751 (PHE, Genbank DQ859059) and the 2016 epidemic Asian lineage isolate PRVABC-59 (PHE, GenBank KX377337). U-87 MG cells were seeded as a monolayer one day before infection. Cells were incubated with ZIKV at 37°C with 5% CO_2_ until the different time points of harvest.

### Quantification of ZIKV RNA using qRT-PCR

Extraction of ZIKV RNA from infected cells was performed using an RNA purification kit following manufacturer recommendations (NucleoSpin RNA II, MACHEREY-NAGEL AG, Düren, Germany). Reverse transcription was done with random primers (Life Technologies, Basel, Switzerland) and SuperScript™ II Reverse Transcriptase (ThermoFisher Scientific, Basel, Switzerland). The quantitative real-time reverse transcription-PCR (qRT-PCR) was performed using a TaqMan probe amplification system to quantify intracellular ZIKV RNA. The primers and TaqMan probes used for qRT-PCR are depicted in Table 1.

**Table 1.**
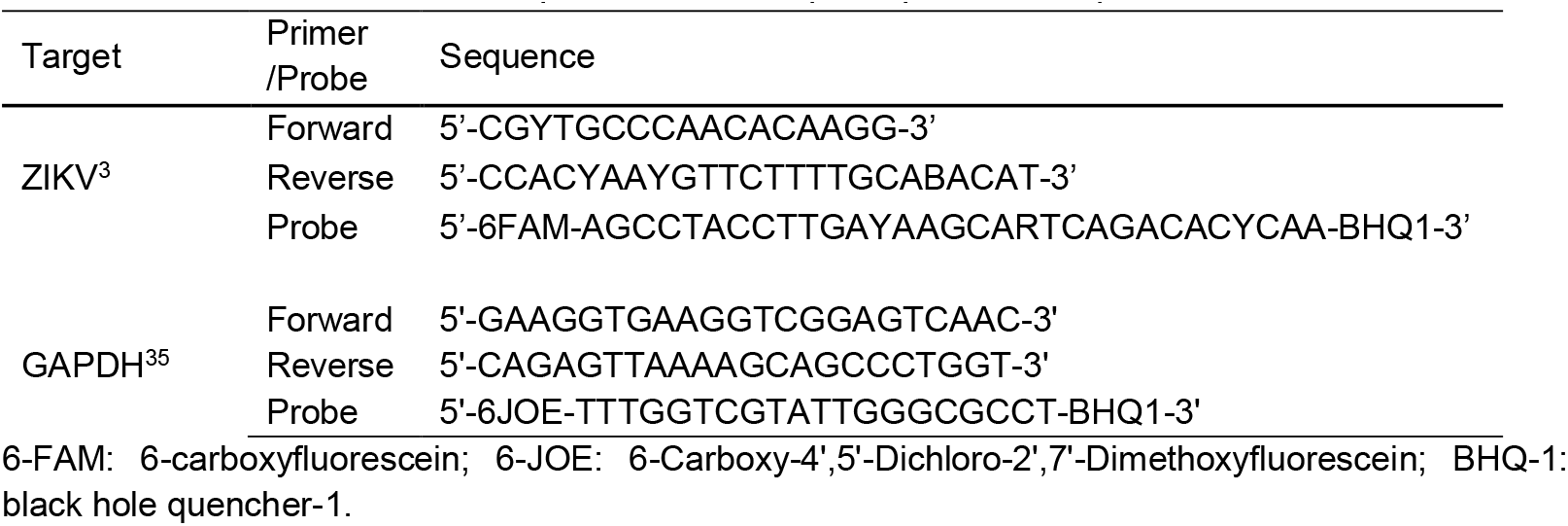
The primers and TaqMan probes for qRT-PCR.

### Plaque assay

Vero cells were seeded at 7×10^5^ cells per well in a 6-well plate (Nunc, Basel, Switzerland) with DMEM medium supplemented with 10% FBS. After 16 to 24 hours, the medium was gently removed, and then 400 μL of a 10-fold serial dilution of virus was added to each well. The plate was incubated at 37°C with 5% CO_2_ for 2 hours. Following incubation, the inoculum was carefully removed, and 3 mL of 1% methyl cellulose (4000 centipoises, Sigma-Aldrich) overlay medium (DMEM supplemented with 2% FBS and 1 mM sodium pyruvate) was added to each well. After 2 days of incubation at 37°C with 5% CO_2_, white plaques became visible to the naked eye. The overlay was removed, and the wells were washed twice with phosphate-buffered saline (PBS, Sigma-Aldrich). Cells were then stained with 1% crystal violet solution for 1 hour. Finally, the plate was rinsed with water, and the white plaques were counted.

### Immunofluorescence microscopy

Cells were seeded on glass coverslips in a 24-well plate (Nunc) in the appropriate culture medium and infected with ZIKV at a multiplicity of infection (MOI) of 5. After incubation, infected cells were fixed with 4% paraformaldehyde (Electron Microscopy Sciences, Pennsylvania, USA) in PBS at room temperature for 15 minutes, followed by permeabilization with 0.1% Triton X-100 in PBS for 10 minutes. ZIKV was detected using a 1:100 dilution of the primary antibody ZKA64 (AG-27B-6004PF, AdipoGen, Lausanne, Switzerland) for 1 hour. Cells were then washed with PBS and incubated with an anti-human IgG-A488 secondary antibody (A10631, Life Technologies) and rhodamine-conjugated concanavalin A (RL-1002, Vector Laboratories, California, USA) for 30 minutes. Afterward, cells were washed with distilled water (ddH_2_O) and mounted on glass slides using DAPI mounting medium (P36941, Life Technologies). Images were captured at 40x magnification using an inverted fluorescence microscope (LSM 880, Zeiss, Oberkochen, Germany).

### Transmission electron microscopy

Cells were seeded in T25 culture flasks (BD Falcon, New Jersey, USA) and infected with ZIKV as described previously at an MOI of 5. At different time points, the growth medium was removed. Cells were subsequently detached with cell scrapers (CLS3010, Corning, New York, USA) and fixed in 2% paraformaldehyde (15714, Electron Microscopy Sciences) and 1.25% glutaraldehyde (G5882, Sigma-Aldrich) in 0.1M phosphate buffer (PB, pH 7.2) for a minimum of 30 minutes at room temperature. Samples were then treated with 1% osmium tetroxide (OsO4, Electron Microscopy Sciences) in ddH_2_O for 1 hour. The samples were then washed three times in ddH_2_O and then dehydrated in a graded ethanol series (50, 70, 80, 90 and 99% for 3 minutes each; 100% for 20 minutes twice). This was followed by infiltration in Epon (EM0300, Sigma-Aldrich) at graded concentrations (Epon 1/3 ethanol-30 minutes; Epon 1/1 ethanol-30 minutes; Epon 3/1 ethanol-30 minutes; Epon 1/1-1 hour; Epon 1/1-16 hours) and finally polymerized for 8 hours at 70°C in oven. Ultrathin sections (70 nm) of resin-embedded cells were prepared using an ultramicrotome (Ultracut S, Leica, Vienna, Austria) and placed on EM grids (G100H-Cu, Electron Microscopy Sciences). Post-staining was performed with 2% uranyl acetate and 0.5% lead citrate to enhance contrast. Images were collected with a transmission electron microscope Philips CM100 (Thermo Fisher Scientific, Waltham, MA USA) at an acceleration voltage of 80kV with a TVIPS TemCam-F416 digital camera (TVIPS GmbH, Gauting, Germany).

### Quantification of convoluted membranes

The ultrastructure of convoluted membranes (CMs) was assessed by randomly selecting 50 cell images obtained through TEM. The percentage of CMs induced by the Asian and African strains was quantified to compare strain-specific effects. Statistical analysis was performed using Student’s t-test.

### Cryo-electron microscopy sample preparation and cryo-focused ion beam scanning electron microscopy

The 200-mesh Quantifoil Au grids (Q2100AR2, Electron Microscopy Sciences) were plasma-cleaned by CCU-010 (Safematic, Zizers, Switzerland) for 120 seconds, placed in a 24-well plate, and incubated in DMEM with 10% FBS for 30 minutes at room temperature. Subsequently, the medium was removed and 8×10^4^ cells were seeded with growth medium on the grids. After 16-24 hours, the grids were incubated with ZIKV at an MOI of 10 at 37°C with 5% CO_2_ for 24 hours. The grids were plunge-frozen into liquid ethane using Vitrobot Mark IV (ThermoFisher Scientific). The blotting chamber was set to 70% humidity, and grids were blotted (blot force of 20) twice, for 5 and 2 seconds, respectively, using Whatman type 1 paper (WHA1001325, Whatman, Massachusetts, USA). Frozen grids were inserted into AutoGrids (ThermoFisher Scientific) for cryo-FIB-SEM milling.

Aquilos 2 Cryo-FIBSEM (ThermoFisher Scientific) was used for lamella milling. The grid was first sputter coated 30 seconds with platinum, followed by organometallic platinum deposition for 90 seconds. The grid was set at 10° milling position and lamella thinning was performed using AutoTEM Cryo software (ThermoFisher Scientific Inc., US) starting at 1 nA to achieve an initial lamella thickness of 2.2 μm. Progressive thinning continued in steps: 1.4 μm thickness at 0.5 nA, 900 nm at 0.3 nA, 500 nm at 0.1 nA and 200 nm at 30 pA. Following cryo-FIB-SEM milling, the grids were transferred to the cryo-grid box and stored in liquid nitrogen.

### Cryo-electron tomography and 3D reconstruction

Data acquisition was performed on a Titan Krios G4 instrument operated at 300 kV equipped with a SelectrisTM X energy filter and a Falcon 4i camera (Thermo Fisher Scientific Inc., US). Tomographic data were collected using the Tomo5 software package (Thermo Fisher Scientific Inc., US). Lamella-overview montages were acquired at a magnification of ×11,500 (pixel size, 2.236 nm). Tilt series were recorded at a magnification of ×53,000, with a pixel size of 2.42 Å and stored in EER file format. Data collection followed a dose-symmetric tilt scheme with 2° angular increments, with 1.6 e−/Å^2^ per tilt and a target defocus of −4.5 μm. The angular range spanned from −70° to 50°, resulting in a total accumulated dose of 100 e−/Å^2^.

Frames were summed to get at least 0.8 e−/px and motion corrected with alignframes command from IMOD package version 5.0.128. Tomogram reconstruction was done with etomo from IMOD package version 5.0.1.

## Results

### African and Asian lineages of ZIKV exhibit comparable replication dynamics

To determine the key time points of the infection process in the human glioblastoma-derived U-87 MG cell line, we investigated the replication dynamics of ZIKV strains MP1751 (African lineage) and PRVABC59 (Asian lineage). Kinetics of viral RNA synthesis and release of infectious viral particles were monitored over time (Figure 1).

**Figure 1.**
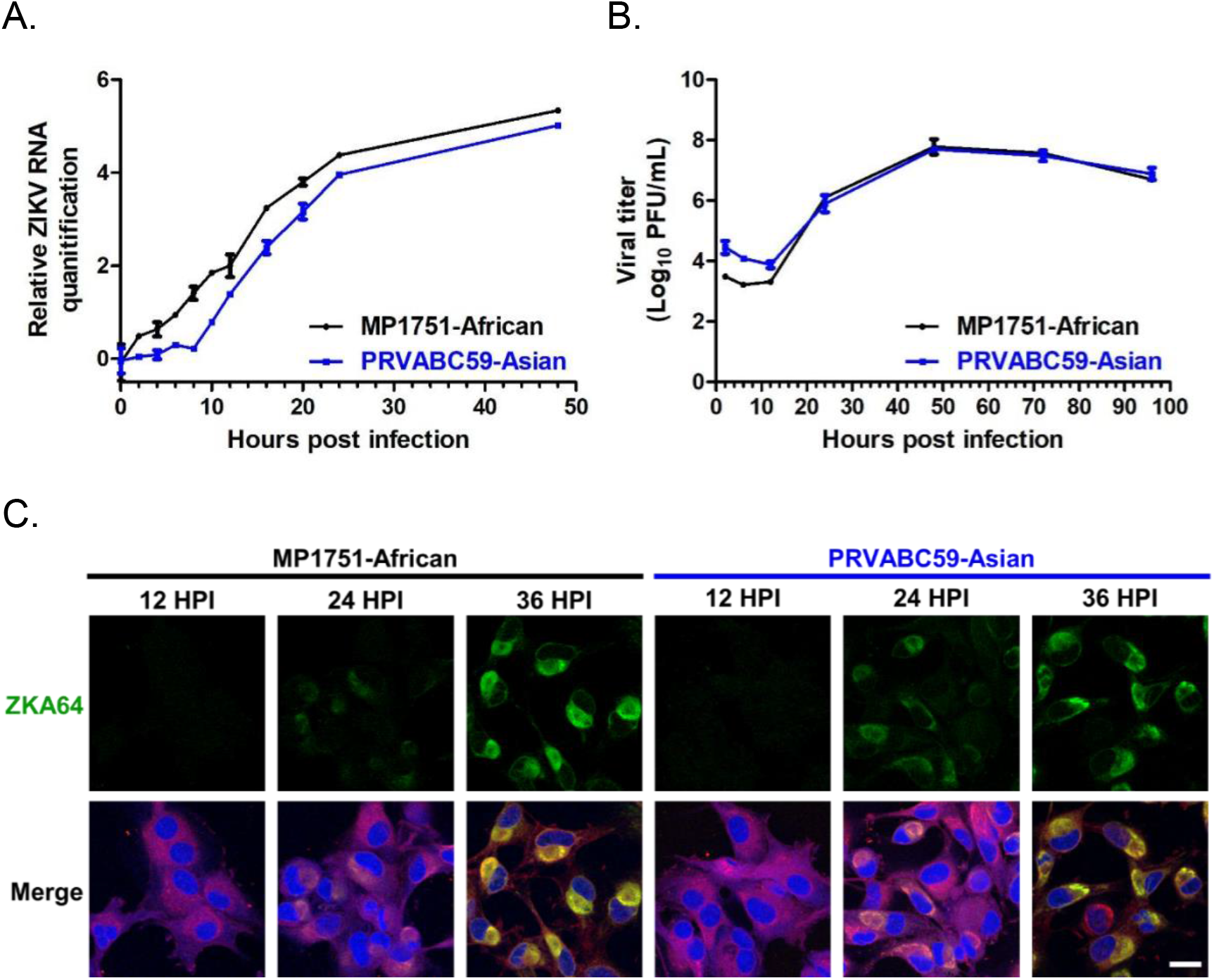
Replication dynamics of ZIKV in U-87 MG cells. U-87 MG cells were infected with African and Asian linage ZIKV at a MOI of 5 PFU/cell. A. Quantification of ZIKV RNA levels over time by RT-qPCR, normalized to GAPDH. Data represent the mean ± SD of three biological replicates. B. Extracellular viral titers measured by plaque assay. Data represent the mean ± SD of duplicate samples. C. Representative temporal expression of ZIKV E glycoprotein by immunofluorescence at 12, 24, and 36 hours post-infection (HPI). ZIKV was detected with the ZKA64 monoclonal antibody (green), cell surfaces were labelled with rhodamine-conjugated concanavalin A (red) and nuclei were counterstained with DAPI (blue). Scale bar represents 20 μm.

In comparison to the MP1751-African strain, which demonstrated early initiation of viral RNA synthesis, the PRVABC59-Asian strain exhibited an ∼8-hour delay before entering the exponential phase of replication (Figure 1A). Despite this initial lag, both strains reached comparable RNA levels at later time points. For both ZIKV strains, infectious virions became detectable in the supernatant at 12 hours post-infection (HPI) and viral titers continued to rise, corresponding to an approximate 4-log increase relative to the initial inoculum (Figure 1B). The progression of viral replication was further supported by confocal microscopy, which revealed initial ZIKV E glycoprotein expression at 24 HPI in both strains, with expression levels increasing by 36 HPI. At later time points, infected cells commonly displayed crescent-shaped nuclei surrounded by viral replication complexes (Figure 1C). By 48 HPI, the majority of infected cells had begun to detach, indicating cytopathic effects associated with late-stage infection (data not shown). Further exploration of these intracellular sites revealed co-localization of viral RNA with the calreticulin endoplasmic reticulum (ER) marker, suggesting a strong spatial association between the replication complexes and ER membranes (Figure 2).

**Figure 2.**
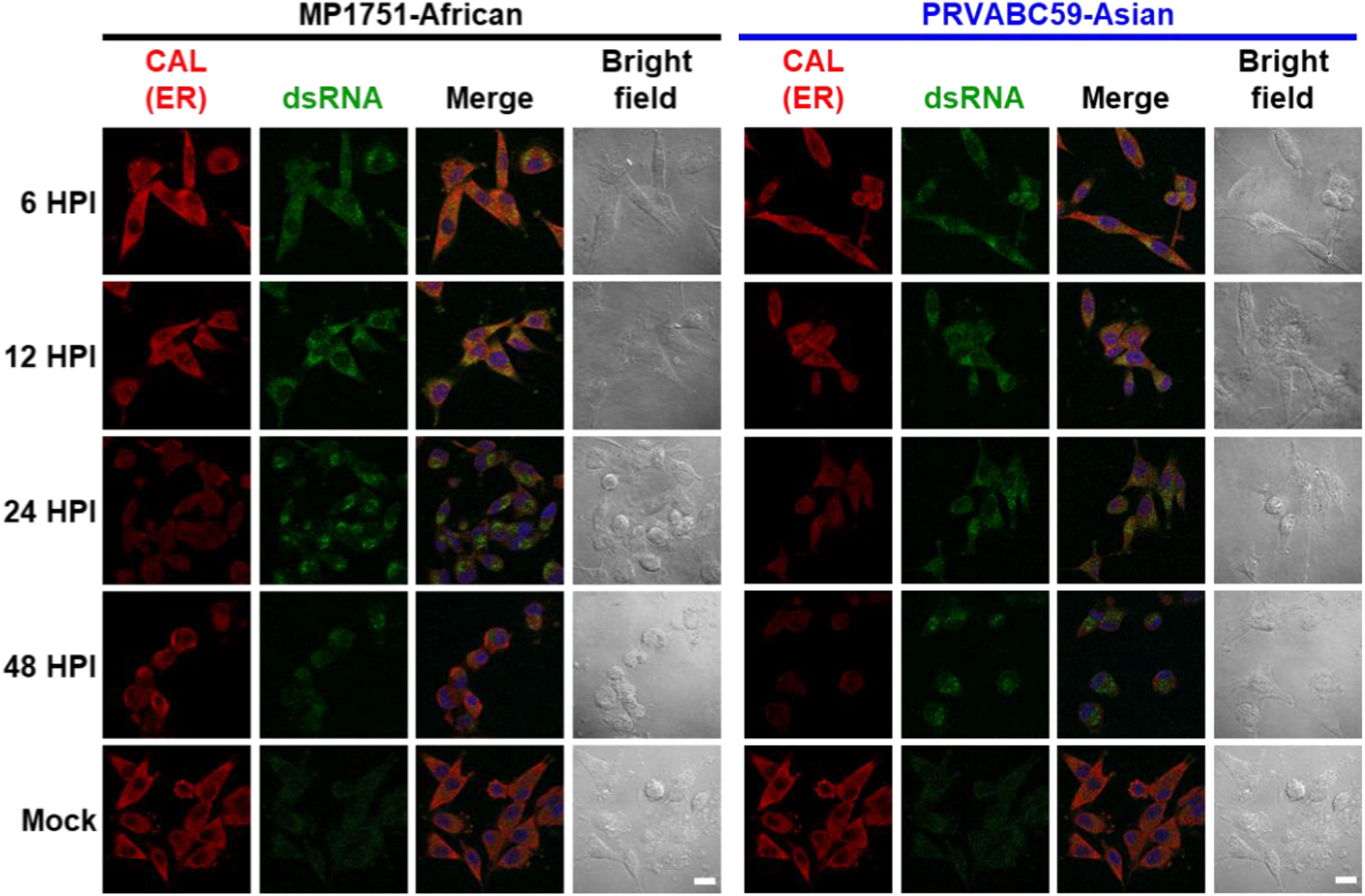
Colocalization of ER and viral dsRNA synthesis sites in ZIKV-infected U-87 MG cells. Cells were infected at an MOI of 5 PFU/cell and analyzed at 6, 12, 24, and 48 HPI. The ER was visualized by immunofluorescence with calreticulin (red), replicating viral dsRNA was detected with an anti-dsRNA antibody (green), and nuclei were counterstained with DAPI (blue). Scale bars represent 20 μm.

Double-stranded RNA, a hallmark of RNA virus replication, became evident by 6 HPI in cells infected with either strain, consistent with the replication kinetics observed by RT-qPCR (Figure 2). Unlike infected cells, mock-treated controls showed no detectable dsRNA, confirming the absence of viral replication. However, they maintained strong calreticulin staining, reflecting an intact ER network (Figure 2).

Multiple viral replication clusters were observed per cell, consistently observed in the perinuclear region. These clusters co-localized with the ER signal and no substantial accumulation or remodelling of the ER membrane was evident at any time point.

### Ultrastructural analysis during early ZIKV replication reveals lineages-specific differences

As the above data indicated that assembled viral particles begin to appear between 12 and 24 HPI, we selected these time points for ultrastructural investigation. Using TEM, we analysed the cytoarchitectural changes in ZIKV-infected U-87 MG cells during this early phase of replication (Figure 3).

**Figure 3.**
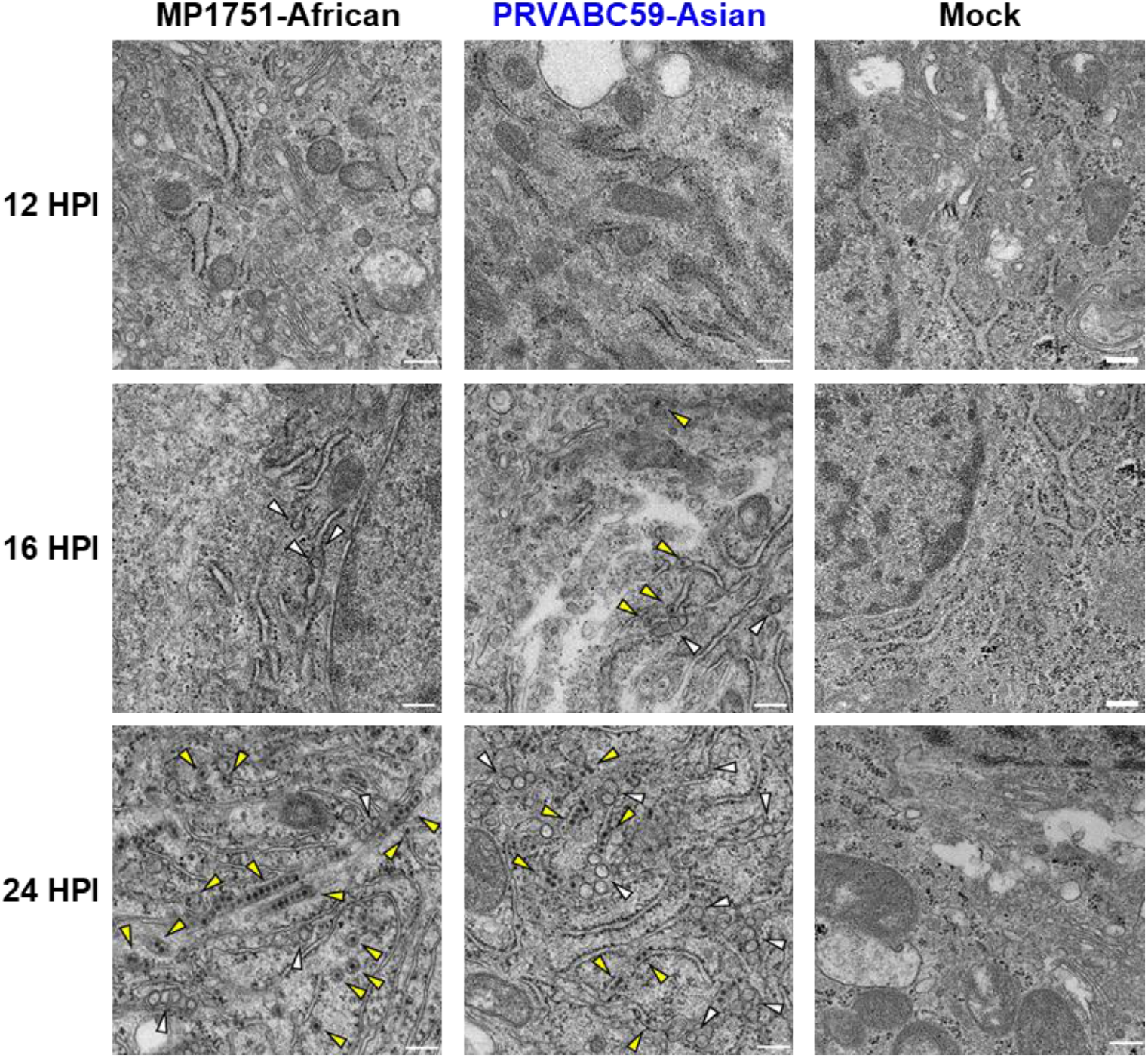
TEM analysis of ZIKV-induced ultrastructural changes in U-87 MG cells. Representative 70-nm sections of resin-embedded U-87 MG cells infected at an MOI of 5 PFU/cell with MP1751-African or PRVABC59-Asian strains, alongside mock-treated controls. Virus-induced vesicles (white arrowheads) and viral particles (yellow arrowheads) are indicated. Scale bars represent 200 nm.

At 12 HPI, no distinct cytoplasmic differences were observed between mock-treated and ZIKV-infected U-87 MG cells (Figure 3). By 16 HPI, ZIKV-induced vesicle formation became apparent and assembled viral particles became visible by 24 HPI. Interestingly, cells infected with the African-lineage strain displayed distinct peapod-like structures containing multiple viral particles at 24 HPI.

Cryo-EM was used to further characterize the cytoarchitecture of ZIKV-infected U-87 MG cells in a near-native state (Figure 4). Consistent with observations from conventional TEM, viral particles were found in associated with the smooth ER (Figure 4A). High-resolution cryo-EM imaging enabled the precise measurement of immature viral particle diameters, which averaged 45.2 ± 5.3 nm for the MP1751-African strain and 45.2 ± 6.2 nm for the PRVABC59-Asian strain (Figure 4B). Notably, virions at different stages of maturation were packaged together in the same compartment in cells infected with the PRVABC59-Asian strain (Figure 4A). In contrast, in cells infected with the MP1751-African strain, virions and vesicles appeared compartmentalized into separate ER structures.

**Figure 4.**
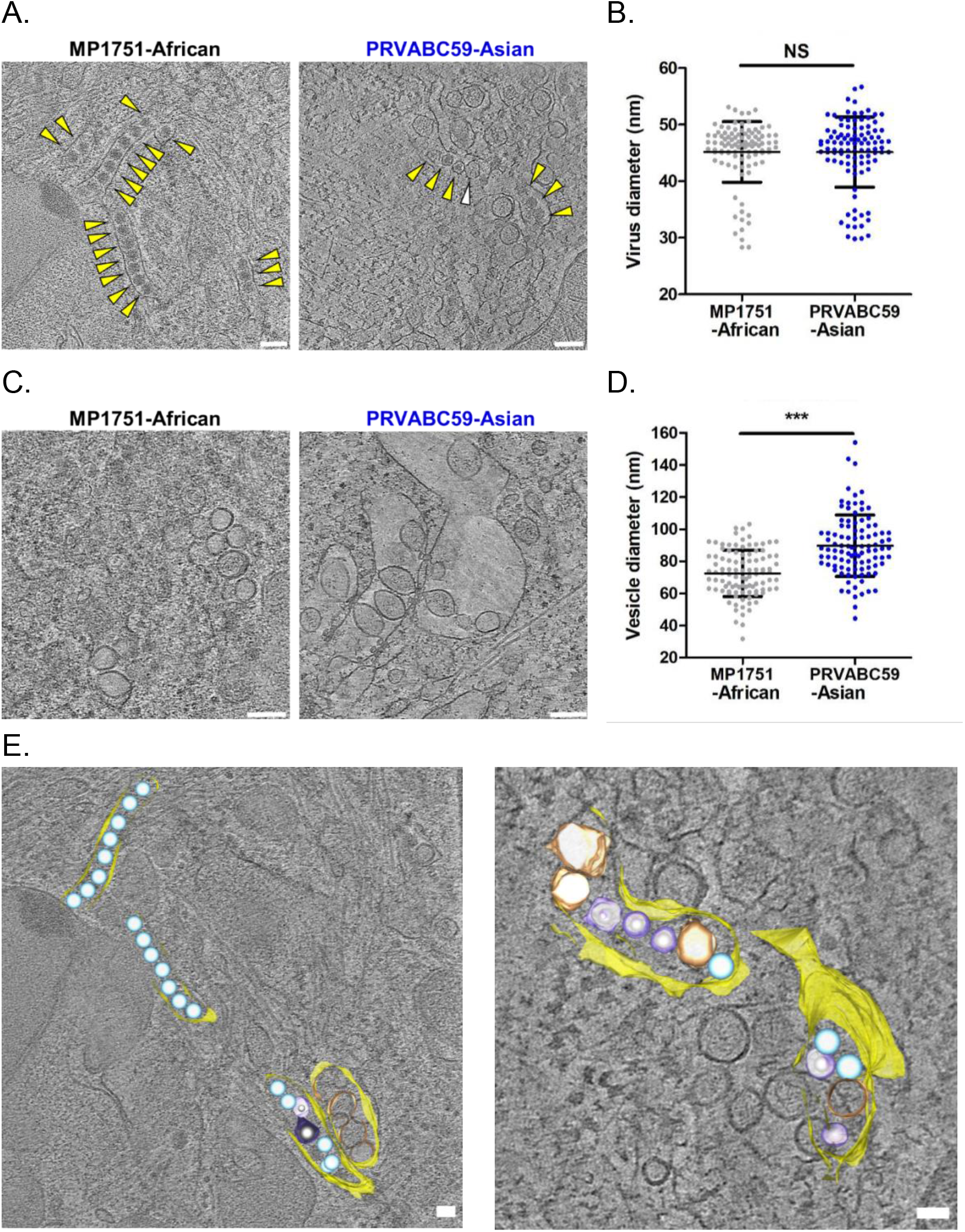
Ultrastructural analysis of ZIKV-infected U-87 MG cells by cryo-ET. A. Tomographic reconstructions of cells infected at an MOI of 10 PFU/cell with MP1751 (left) or PRVABC59 (right) at 24 HPI. White arrowheads indicate virus-induced vesicles; yellow arrowheads mark viral particles. Scale bars represent 200 nm. B. Viral particle diameters (n = 100 per strain) presented as mean ± SD. C. Representative cryo-ET slices showing vesicles and double-membrane structures in MP1751- and PRVABC59-infected cells. Scale bars represent 200 nm. D. Vesicle diameters quantified for each strain (n = 100 vesicles; mean ± SD). Statistical comparisons by unpaired t-test: NS = not significant; ***p < 0.001. E. 3D surface renderings of infected cells: virus (light blue), ER (yellow), vesicles (golden brown), viral envelope (purple), nucleocapsid (white). Scale bars represent 100 nm.

Additionally, the spacing between the ER membrane and individual virions was consistently uniform in MP1751-African-infected cells. The organized disposition of viral particles in MP1751-African-infected cells, was not present in PRVABC59-Asian-infected cells, which displayed greater morphological heterogeneity in the appearance of virus-induced vesicles (Figure 4C). These observations are consistent with the structural differences previously noted using conventional TEM (Figure 3). We measured the diameter of virus-induced vesicles, by randomly selecting 100 vesicles (Figure 4D). Vesicles induced by the PRVABC59-Asian strain were significantly larger (89.7 ± 19.2 nm) than those in MP1751-African-infected cells (72.5 ± 14.4 nm).

To further investigate the replication sites, we performed cryo-ET at 24 HPI (Figure 4E, Supplementary files S1 and S2). We observed multiple contact points between double-membrane vesicles and nearby viral particles, although membrane fusion between vesicles could not be detected.

### Asian lineage ZIKV induces more extensive membrane rearrangement than the African lineage

TEM analysis of ZIKV-infected U-87 MG cells revealed abundant virions and virus-induced vesicles throughout the cytoplasm at 48 HPI (Figure 5A and B). In this late stage, the ER underwent pronounced reorganization, with the appearance of convoluted membranes (CMs). These structures were present as dense, highly irregular, and tightly packed membranous structures, typically localized in the perinuclear region and closely associated with ER-derived membranes (Figure 5A and B). Quantification showed a significantly higher number of CMs in PRVABC59-infected cells compared to MP1751-infected cells (Figure 5C), demonstrating lineage-specific differences in membrane remodelling during replication.

**Figure 5.**
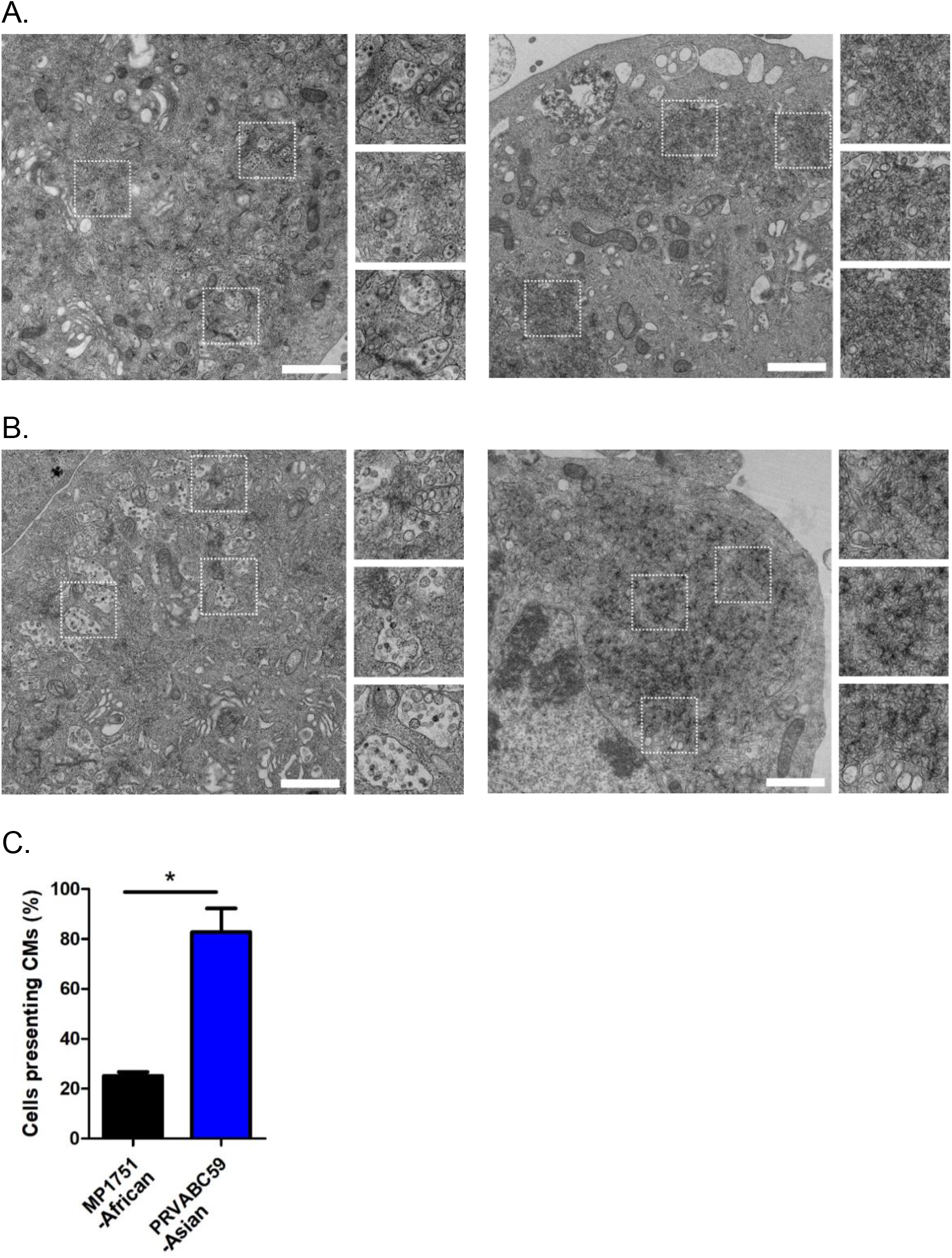
ZIKV-induced rearrangement of intracellular membranes forming convoluted membranes (CMs). TEM of 70-nm resin-embedded sections of U-87 MG cells infected at an MOI of 5 PFU/cell with (A) MP1751 and (B) PRVABC59 strains, imaged at 48 HPI. Details of highlighted structures in the infected cells are shown on the right panels of each figure. Scale bar represents 1 µm. B. Percentage of cells containing CMs structures in 50 randomly selected infected cells per strain. Data are shown as mean ± SD; *p < 0.05.

## Discussion

This study presents a high-resolution ultrastructural comparison of ZIKV replication in U-87 MG cells by combining conventional TEM, cryo-FIB-SEM milling and cryo-ET to compare two phylogenetically distinct strains: African-lineage MP1751 and Asian-lineage PRVABC59. The results underscore both conserved and lineage-specific features of ZIKV replication and highlight the distinct cellular remodelling strategies employed by each lineage. Notably, these two lineages are also associated with divergent pregnancy outcomes: clinically, infections with the Asian-lineage strain (PRVABC59) have been more frequently associated with congenital Zika syndrome, while reports of adverse pregnancy outcomes linked to African-lineage strains in humans remain limited^25,9,26^.

Consistent with prior studies on flavivirus replication, both ZIKV lineages reorganized the ER into replication complexes composed of vesicle packets and virus-induced compartments^18,19,14^. During the early stages of infection, MP1751 displayed rapid initiation of viral RNA synthesis and vesicle formation, with viral particles becoming detectable as early as 16 HPI. In contrast, PRVABC59 replication showed an 8-hour lag. Despite these kinetic differences, both strains achieve similar RNA levels and infectious titers at later time points. Immunofluorescence analyses confirmed that replication clusters localized in the perinuclear region, colocalising with ER markers. Interestingly, while MP1751-infected cells showed an organized architecture with early appearance of peapod-like virion arrangements, PRVABC59-infected cells exhibited more heterogeneous replication sites, containing virions at various maturation stages within the same vesicle. Cryo-ET confirmed this observation and further demonstrated that in PRVABC59-infected cells, vesicles and virions were often packaged together within the same compartment, whereas MP1751 maintained separate organization. Although mean virion sizes were comparable (about 45 nm), induced vesicle sizes differ significantly, with PRVABC59 vesicles measuring 89.7 ± 19.2 nm and MP1751 vesicles 72.5 ± 14.4 nm. These data align with previous studies in other cell types showing that Asian-lineage ZIKV tends to induce larger vesicles than its African-lineage counterparts^14^. At later stages of infection (48 HPI), pronounced ER remodelling produced CMs predominantly in PRVABC59-infected cells. These dense, irregular, perinuclear structures were significantly higher in cells infected with the Asian lineage compared to the African lineage strain and may reflect therefore lineage-specific effect of viral replication. Although the precise function of CMs remains elusive, previous studies on other flaviviruses have proposed a roles in polyprotein processing^20^, storage of replication intermediates^27^, or immune evasion^19,28^. The significant increase in CMs in PRVABC59-infected cells may point to higher replication stress, enhanced replication demands, or alternative mechanisms of host manipulation. CMs have also been reported in other virus-infected models. Several studies have highlighted these structures, not only in flaviviruses^20,21,29–31^ but also in other positive-strand RNA viruses such as coronaviruses^32–34^. Also, both Picornaviridae and Arteriviridae are capable of inducing CM–like structures or highly complex ER-derived membrane networks^35,36^. Although the exact function of these structures remains unclear, their close proximity to the viral replication complexes suggests they may play a role in promoting viral replication.

Methodologically, we demonstrate that cryo-ET preserves native membrane architecture far better than chemically fixed TEM, although artifacts such as ice crystallization and sample compression can limit resolution at vesicle–ER interfaces. Future use of high-pressure freezing could further mitigate these issues and improve the resolution of vesicle-ER junctions.

In conclusion, our findings reveal both conserved and lineage-specific ultrastructural features of ZIKV replication, with Asian lineage driving more extensive membrane remodelling and CM formation than African lineage. These differences may underlie variations in pathogenic potential and warrant further investigations in more physiologically relevant models, such as brain organoids, to elucidate how early replication events and membrane remodelling contribute to neurotropism and congenital disease.

## References

1. WHO statement on the first meeting of the International Health Regulations (2005) (IHR 2005) Emergency Committee on Zika virus and observed increase in neurological disorders and neonatal malformations. https://www.who.int/news/item/01-02-2016-who-statement-on-the-first-meeting-of-the-international-health-regulations-(2005)-(ihr-2005)-emergency-committee-on-zika-virus-and-observed-increase-in-neurological-disorders-and-neonatal-malformations.

2. Duong, V., Dussart, P. & Buchy, P. Zika virus in Asia. Int. J. Infect. Dis. 54, 121–128 (2017).

3. Lanciotti, R. S. et al. Genetic and Serologic Properties of Zika Virus Associated with an Epidemic, Yap State, Micronesia, 2007 - Volume 14, Number 8—August 2008 - Emerging Infectious Diseases journal - CDC. doi:10.3201/eid1408.080287.

4. Cao-Lormeau, V.-M. et al. Zika virus, French polynesia, South pacific, 2013. Emerg. Infect. Dis. 20, 1085–1086 (2014).

5. de Oliveira, W. K. et al. Zika Virus Infection and Associated Neurologic Disorders in Brazil. N. Engl. J. Med. 376, 1591–1593 (2017).

6. Cugola, F. R. et al. The Brazilian Zika virus strain causes birth defects in experimental models. Nature 534, 267–271 (2016).

7. Haddow, A. D. et al. Genetic Characterization of Zika Virus Strains: Geographic Expansion of the Asian Lineage. PLoS Negl. Trop. Dis. 6, e1477 (2012).

8. Song, B.-H., Yun, S.-I., Woolley, M. & Lee, Y.-M. Zika virus: History, epidemiology, transmission, and clinical presentation. J. Neuroimmunol. 308, 50–64 (2017).

9. Beaver, J. T., Lelutiu, N., Habib, R. & Skountzou, I. Evolution of Two Major Zika Virus Lineages: Implications for Pathology, Immune Response, and Vaccine Development. Front. Immunol. 9, 1640 (2018).

10. Hu, T., Li, J., Carr, M. J., Duchêne, S. & Shi, W. The Asian Lineage of Zika Virus: Transmission and Evolution in Asia and the Americas. Virol. Sin. 34, 1–8 (2019).

11. Darmuzey, M. et al. Epidemic Zika virus strains from the Asian lineage induce an attenuated fetal brain pathogenicity. Nat. Commun. 15, 10870 (2024).

12. Sirohi, D. & Kuhn, R. J. Zika Virus Structure, Maturation, and Receptors. J. Infect. Dis. 216, S935–S944 (2017).

13. Offerdahl, D. K., Dorward, D. W., Hansen, B. T. & Bloom, M. E. Cytoarchitecture of Zika virus infection in human neuroblastoma and Aedes albopictus cell lines. Virology 501, 54–62 (2017).

14. Cortese, M. et al. Ultrastructural Characterization of Zika Virus Replication Factories. Cell Rep. 18, 2113–2123 (2017).

15. Ci, Y. & Shi, L. Compartmentalized replication organelle of flavivirus at the ER and the factors involved. Cell. Mol. Life Sci. 78, 4939–4954 (2021).

16. Welsch, S. et al. Composition and Three-Dimensional Architecture of the Dengue Virus Replication and Assembly Sites. Cell Host Microbe 5, 365–375 (2009).

17. Gillespie, L. K., Hoenen, A., Morgan, G. & Mackenzie, J. M. The Endoplasmic Reticulum Provides the Membrane Platform for Biogenesis of the Flavivirus Replication Complex. J. Virol. 84, 10438–10447 (2010).

18. Westaway, E. G., Mackenzie, J. M., Kenney, M. T., Jones, M. K. & Khromykh, A. A. Ultrastructure of Kunjin virus-infected cells: colocalization of NS1 and NS3 with double-stranded RNA, and of NS2B with NS3, in virus-induced membrane structures. J. Virol. 71, 6650–6661 (1997).

19. Junjhon, J. et al. Ultrastructural Characterization and Three-Dimensional Architecture of Replication Sites in Dengue Virus-Infected Mosquito Cells. J. Virol. 88, 4687–4697 (2014).

20. Sirohi, D. et al. The 3.8 Å resolution cryo-EM structure of Zika virus. Science 352, 467– 470 (2016).

21. Prasad, V. M. et al. Structure of the immature Zika virus at 9 Å resolution. Nat. Struct. Mol. Biol. 24, 184–186 (2017).

22. Sevvana, M. et al. Refinement and Analysis of the Mature Zika Virus Cryo-EM Structure at 3.1 Å Resolution. Struct. Lond. Engl. 1993 26, 1169-1177.e3 (2018).

23. Winey, M., Meehl, J. B., O’Toole, E. T. & Thomas H Giddings, J. Conventional transmission electron microscopy. Mol. Biol. Cell 25, 319 (2014).

24. Al-Amoudi, A. et al. Cryo-electron microscopy of vitreous sections. EMBO J. 23, 3583– 3588 (2004).

25. Al-Amoudi, A., Norlen, L. P. O. & Dubochet, J. Cryo-electron microscopy of vitreous sections of native biological cells and tissues. J. Struct. Biol. 148, 131–135 (2004).

26. Ravi, R. T., Leung, M. R. & Zeev-Ben-Mordehai, T. Looking back and looking forward: contributions of electron microscopy to the structural cell biology of gametes and fertilization. Open Biol. 10, 200186 (2020).

27. McCafferty, C. L. et al. Integrating cellular electron microscopy with multimodal data to explore biology across space and time. Cell 187, 563–584 (2024).

28. Keller, J. & Fernández-Busnadiego, R. In situ studies of membrane biology by cryo-electron tomography. Curr. Opin. Cell Biol. 88, 102363 (2024).

29. Mahamid, J. et al. Visualizing the molecular sociology at the HeLa cell nuclear periphery. Science 351, 969–972 (2016).

30. Turk, M. & Baumeister, W. The promise and the challenges of cryo-electron tomography. FEBS Lett. 594, 3243–3261 (2020).

31. Rigort, A. et al. Focused ion beam micromachining of eukaryotic cells for cryoelectron tomography. Proc. Natl. Acad. Sci. 109, 4449–4454 (2012).

32. Buckley, G. et al. Automated cryo-lamella preparation for high-throughput in-situ structural biology. J. Struct. Biol. 210, 107488 (2020).

33. Berger, C. et al. Cryo-electron tomography on focused ion beam lamellae transforms structural cell biology. Nat. Methods 20, 499–511 (2023).

34. Dubochet, J. & Sartori Blanc, N. The cell in absence of aggregation artifacts. Micron Oxf. Engl. 1993 32, 91–99 (2001).

35. Shirato, K., Kawase, M. & Matsuyama, S. Middle East Respiratory Syndrome Coronavirus Infection Mediated by the Transmembrane Serine Protease TMPRSS2. J. Virol. 87, 12552–12561 (2013).

36. Romero-Brey, I. et al. Three-Dimensional Architecture and Biogenesis of Membrane Structures Associated with Hepatitis C Virus Replication. PLOS Pathog. 8, e1003056 (2012).

37. Hartenian, E. et al. The molecular virology of coronaviruses. J. Biol. Chem. 295, 12910–12934 (2020).

38. Miorin, L. et al. Three-dimensional architecture of tick-borne encephalitis virus replication sites and trafficking of the replicated RNA. J. Virol. 87, 6469–6481 (2013).

39. Arakawa, M. & Morita, E. Flavivirus Replication Organelle Biogenesis in the Endoplasmic Reticulum: Comparison with Other Single-Stranded Positive-Sense RNA Viruses. Int. J. Mol. Sci. 20, 2336 (2019).

40. Garcia, M. D. do N. et al. In Vitro System for Studying Ilhéus Virus, a Neglected Arbovirus: Ultrastructural Characterization of Cytopathology, Morphology, and Morphogenesis. Viruses 17, 320 (2025).

41. Ulasli, M., Verheije, M. H., de Haan, C. A. M. & Reggiori, F. Qualitative and quantitative ultrastructural analysis of the membrane rearrangements induced by coronavirus. Cell. Microbiol. 12, 844–861 (2010).

42. Blanchard, E. & Roingeard, P. Virus-induced double-membrane vesicles. Cell. Microbiol. 17, 45–50 (2015).

43. Snijder, E. J. et al. A unifying structural and functional model of the coronavirus replication organelle: Tracking down RNA synthesis. PLOS Biol. 18, e3000715 (2020).

44. Belov, G. A. et al. Complex Dynamic Development of Poliovirus Membranous Replication Complexes. J. Virol. 86, 302–312 (2012).

45. Junjhon, J. et al. Ultrastructural Characterization and Three-Dimensional Architecture of Replication Sites in Dengue Virus-Infected Mosquito Cells. J. Virol. 88, 4687–4697 (2014).

